# Leptin and adiponectin correlations with body composition and lipid profile in children with Autism Spectrum Disorder

**DOI:** 10.1101/621003

**Authors:** Kamila Castro, Larissa Slongo Faccioli, Ingrid Schweigert Perry, Rudimar dos Santos Riesgo

## Abstract

Leptin and adiponectin have effects on the regulation of appetite and body composition, but evidence of these relationships in children is still limited. Even though investigations of their role in children with ASD are incipient, the nutritional aspects and eating difficulties that these patients may present are increasingly highlighted, often leading to inadequate nutritional status. This cross-sectional controlled study investigated the levels of adipokines in ASD children in comparison with healthy controls, and their correlations with nutritional aspects and lipid profile. A total of 80 participants (40 ASD and 40 controls) were included and evaluated through anthropometric variables, body composition, and blood samples. ASD participants showed higher levels of leptin, no changes of adiponectin levels in comparison with typically developing children, and a positive correlation between leptin and fat mass. This novel finding supports the role of leptin as a marker of adiposity in ASD children, which is reiterated by the higher leptin/adiponectin ratio and its correlation with fat mass in patients. Inverse correlation of leptin with HDL-cholesterol could only in certain cases be related to the higher adiposity in patients when compared to controls. These results highlight also the importance of assessing the nutritional status of this population.

## 1. Introduction

Autism spectrum disorder (ASD) affects approximately 1% of US children (“Prevalence of autism spectrum disorders--Autism and Developmental Disabilities Monitoring Network, 14 sites, United States, 2008,” 2012), is a neurodevelopmental disorder characterized by restricted, stereotyped behaviors and impairments in social communication (DSM-V, 2013). In the developing of the disease, a role has been hypothesized for environmental factors, immune dysfunctions, likewise for alterations of neurotransmitter systems (Pardo, 2017).

There are recent reports in the literature about different cytokine levels in children with ASD in comparison with typically developing children. Increased plasma level of interleukins (IL-1β, IL-6, IL-8, and IL-12) and correlations between cytokine levels and impaired verbal communication or aberrant behaviors were described in this population (Ashwood et al., 2011). In individuals without ASD, studies have investigated the correlation between obesity and inflammatory cytokines, such as adiponectin - an anti-inflammatory cytokine. In adolescent girls with central obesity, a significant decrease in the anti-inflammatory adiponectin and an increase in the inflammatory leptin values were described, as well as positive correlations between waist to hip ratio and leptin, TNF-α, IL-1β, IL-4 and IL-5 and negative ones with adiponectin (El-Wakkad, Hassan, Sibaii, & El-Zayat, 2013).

Adipokines are hormones synthesized mainly by the adipocytes of the white adipose tissue (Pan & Kastin, 2007). Initially, they were associated with eating disorders and diabetes, with later studies showing the important role in the regulation of immune responses and inflammation. Decreased serum levels of adiponectin were described in a group of ASD patients and the involvement of adiponectin in the pathophysiology of autism was hypothesized (Ouchi, Shibata, & Walsh, 2006).

Leptin is a hormone develop especially in the adipose tissue and in small amounts in the stomach, mammary epithelium, placenta, and heart. The presence of leptin receptors in specific regions of the brain illustrates their potential for being involved in multiple mechanisms related to brain function and structure (Roubos, Dahmen, Kozicz, & Xu, 2012). A recent review indicates that leptin plays roles in immunity, regulation of insulin secretion, sex hormone release, performs lipolysis in adipocytes and modulates plasticity in learning and memory-based behavioral tasks (Van Doorn, Macht, Grillo, & Reagan, 2017). This hormone has an important role in the regulation of food intake and body weight (Klok, Jakobsdottir, & Drent, 2007) and its expression by adipose tissue is also influenced by feeding behavior. Ambroszkiewicz *et al*. (2017) (Ambroszkiewicz et al., 2017) demonstrated that leptin levels were significantly lower in thin children (1.33; 0.65 – 1.62) than in normal weight children (3.06; 1.60 – 5.18). This same study says that leptin emerges as a marker of the degree of adiposity in the young population.

Few studies described that leptin in ASD subjects is higher than in typically developing controls (Ashwood et al., 2008; Blardi et al., 2010; Rodrigues et al., 2014). Also, long-term higher plasma leptin levels in Rett syndrome was described (Blardi et al., 2009). In addition, data on adiponectin levels in these patients are controversial, with reports of higher (Fujita-Shimizu, 2010) or unaltered (Rodrigues, 2014) levels, but correlated with clinical symptoms in both studies (Fujita-Shimizu, 2010; Rodrigues, 2014). Still, reports seeking the association of these adipokines and nutritional status are very scarce in this population, focusing exclusively on body mass index measures, without finding significant associations (Blardi et al., 2010; Rodrigues et al., 2014). An altered plasma lipid profile was also described in these patients (Kim, Neggers, Shin, Kim, & Kim, 2010). We investigated the involvement of the adipokines (leptin and adiponectin) in ASD’ patients in comparison with healthy controls, and their correlations with nutritional aspects and lipid profile.

## 2. Methods

This cross-sectional controlled study was developed with male ASD patients, diagnosed by DSM-5 (DSM-V, 2013) and matched controls by age and weight. Patients were recruited in the Neuropediatric Department at the Hospital de Clínicas de Porto Alegre (HCPA), Brazil, and controls were children and adolescents in continuous follow-up at the Pediatric Service from the same hospital. For both groups, inclusion criteria were age between 3-10 years – this age interval was selected in order to avoid biases from the puberty period (Souza et al., 2012). Chronic use of any medication was an exclusion criterion for both patients and controls. Only patients without any genetic alteration were included in this study.

### 2.1 Clinical variables

Clinic data were accessed by the patients’ medical records. In addition, the scores for the Autism Screening Questionnaire (ASQ) (Rutter, 1996; Sato et al., 2009) and Childhood Autism Rating Scale (CARS) (Pereira, Riesgo, & Wagner, 2008; Rutter & Schopler, 1992) were obtained.

### 2.2 Anthropometrics and Body composition variables

Anthropometric variables [height (cm), weight (kg) and waist circumference (WC)], and body composition variables [fat mass (FM) and fat-free mass (FFM)] were performed according to previously described protocol (Castro et al., 2017). Anthropometrics were done using a wall-mounted stadiometer (Harpenden, Holtain^®^, Crymych, UK) for height, a digital platform scale for weight (Toledo^®^, Model 2096PP/2, São Paulo, Brazil), and a Cescorf^®^ inelastic measuring tape for WC. Body mass index (BMI) was calculated and classified by z –score according to Anthro Plus software (WHO, 2009), and WC was classified according to Taylor et al (2000) (R. W. Taylor, Jones, Williams, & Goulding, 2000). Body composition measurements were performed using a bioelectrical impedance analysis (BIA) device (Biodynamics 450^®^ version 5.1, Biodynamics Corporation, Seattle, WA, USA) and Resting ECG tab electrodes (Conmed Corporation, Utica, NY, USA).

### 2.3 Biochemical variables

Blood samples (6mL) were withdrawn after an overnight fasting. The blood was then centrifuged at 3000rpm for 10 min. Plasma was collected and stored at –80° C until the analysis was done.

Plasma levels of adiponectin and leptin were measured using commercially available kits (Human Leptin Enzyme Immunoassay, Merck, Cat. #A05174 and Human Adiponectin ELISA, Merck, Cat. # EZHADP-61K). The minimum detectable dose of leptin was 7.8 pg/mL and that of adiponectin was 0.891 μg/mL.

Additionally, part of the collected sample was sent to the biochemistry unit of the hospital for analysis of lipid profile for total cholesterol (total-chol), high-density lipoprotein-cholesterol (HDL-chol) and low-density lipoprotein-cholesterol (LDL-chol). The total and LDL cholesterol levels were classified according to the American Academy of Pediatrics (AAP).

### 2.4 Statistical aspects

Statistical Package for Social Sciences 22.0 (SPSS Inc., Chicago, IL) was used. Data were described using absolute and relative frequencies. Shapiro-Wilk statistical test was performed to verify the normality of the variables. Continuous variables were expressed as a mean ± standard deviation and compared through the paired t-test. In addition, the Spearman’s rank correlation coefficient was performed to test correlations between leptin, adiponectin levels and leptin/adiponectin ratio (L/A ratio) and other variables. The level of significance was set at0.05.

### 2.5 Ethical aspects

The study has been approved by the Research Ethics Committee of HCPA (protocol number 16-0464) and was conducted according to the Declaration of Helsinki guidelines. The parents of all the children provided written informed consent.

## 3. Results

The total sample had 80 male participants (40 controls and 40 cases). There was no difference between patients and controls for age (7.8±2.2, 7.7±2.3 years, p=0.891). ASD group presented mean scores 34.84±6.23 for CARS and 22.07±2.73 for ASQ scores.

The anthropometric variables were described in Table 1. There was no difference for weight, height and BMI z-scores per age between controls and cases. The classification for the WC was similar between groups, with 16 patients and 18 controls presenting high values (L. Taylor, Swerdfeger, & Eslick, 2014).

**Table 1.** Anthropometric measurements and body composition variables^†^

| Variables | Controls<br>mean±SD | Cases<br>mean±SD | p |
| --- | --- | --- | --- |
| *Anthropometric* |  |  |  |
| Weight (kg) | 32.5±16.54 | 33.8±14.6 | 0.807 |
| Height (cm) | 128.7±14.5 | 129.2±15.5 | 0.923 |
| BMI (kg/m<sup>2</sup>) | 18.6±4.5 | 19.3±4.3 | 0.627 |
| WC (cm) | 66.1±17.4 | 63.7±15.9 | 0.677 |
| AC (cm) | 21.19±4.85 | 22.32±5.60 | 0.772 |
| *z- score per age* |  |  |  |
| Weight | 1.41±1.3 | 1.07±1.4 | 0.654 |
| Height | 0.62±1.0 | 0.67±1.3 | 0.872 |
| BMI | 1.0±1.4 | 1.4±1.3 | 0.435 |
| *Body composition* |  |  |  |
| FM (kg) | 17.9±7.9 | 26.4±8.5 | 0.000* |
| FFM (kg) | 25.8±10.8 | 22.0±7.9 | 0.005* |

**AC:** Arm circumference; **BMI:** Body mass index; **FFM:** Fat free mass; **FM:** Fat mass; **SD:** standard deviation; **WC:** Waist circumference.
<sup>†</sup>Paired t-test; \*Significant p-values.

The body composition analyzed through BIA showed a significant difference for FM and FFM, ASD group presented higher values for FM (kg) and lower values for FFM (kg) compared to controls (Table 1).

The lipid profile did not demonstrate significant difference between controls and patients for HDL-chol (mg/dl) (52.9±11.3, 57.3±8.4, p=0.243), LDL-chol (mg/dl) (105.8±45.1, 99.6±23.1, p= 0.602) and total cholesterol (mg/dl) (153.3±33.1, 156.9±21.5, p=0.697). Table 2 shows the classified results for total and LDL-chol for both groups. The levels of leptin were significantly different between groups, ASD patients present higher levels compared to controls (1.2±0.5ng/mL, 0.6±0.4ng/mL, p=0.034, respectively) (Figure 1A). There was no difference between groups for adiponectin (Figure 1B), however the leptin/adiponectin ratio (L/A ratio) was higher in ASD patients (Figure 1C).

**Table 2.**
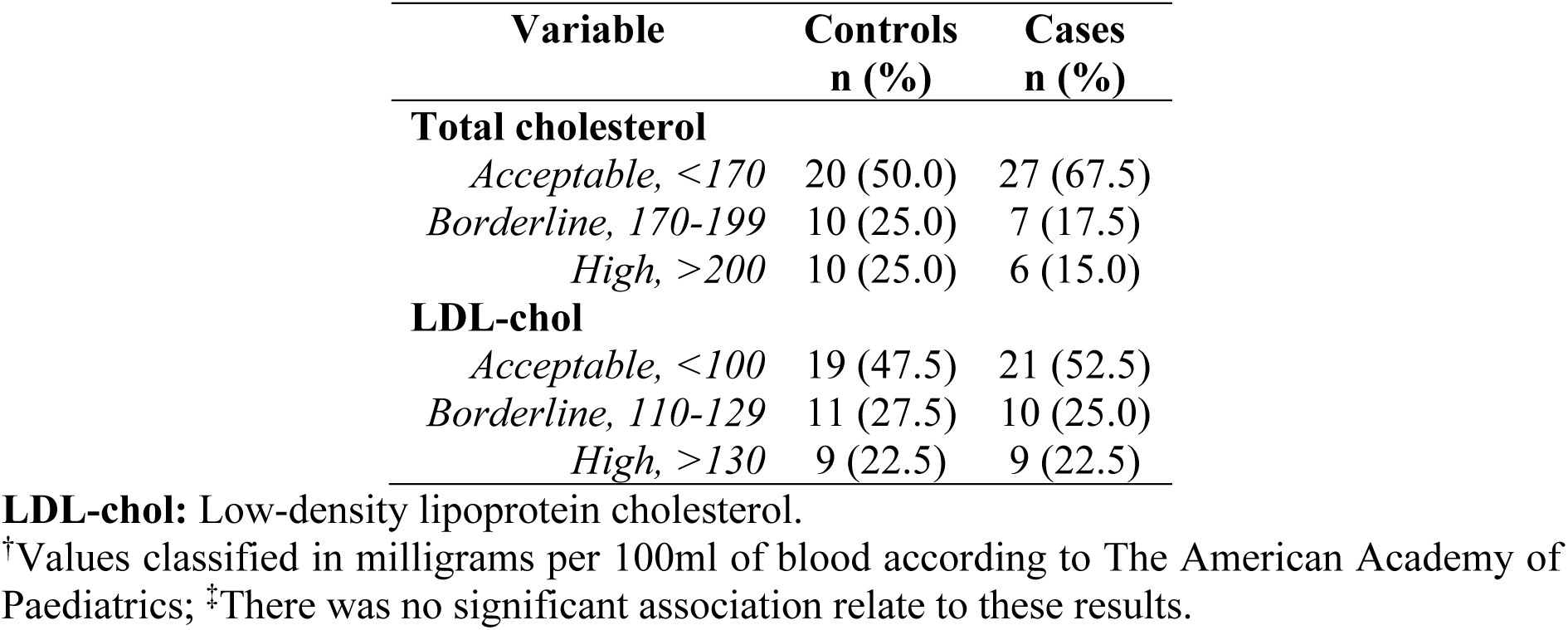
Lipid Profile Classification^†,‡^

**Figure 1.**
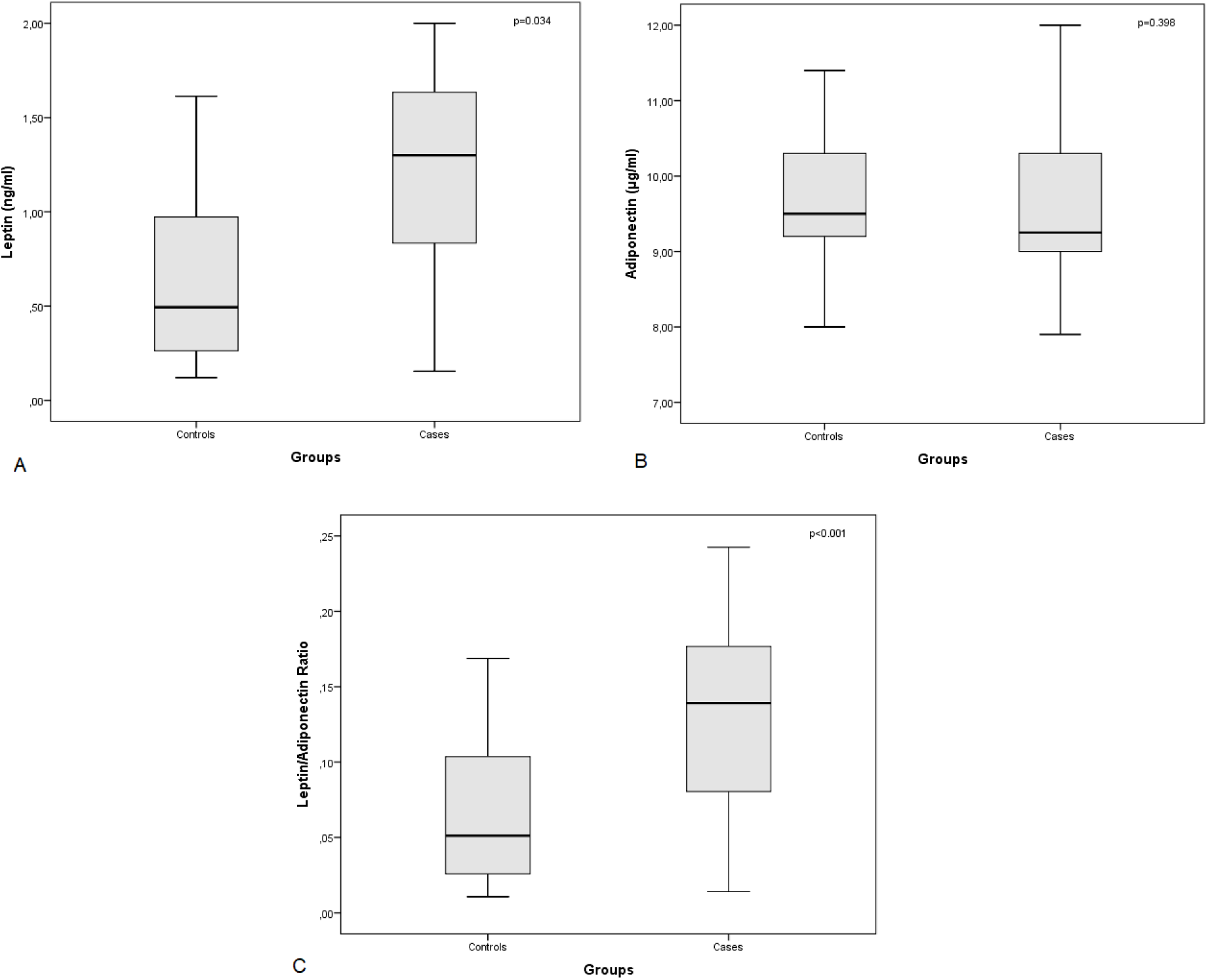
Levels of Leptin, adiponectin and leptin/adiponectin ratio between control and case group. **(A)**Leptin levels between controls (0.6±0.4) and cases (1.2±0.5), paired t-test, p=0.034; **(B)**Adiponectin levels between controls (9.7±0.9) and cases (9.5±0.9), paired t-test, p=0.398; **(C)**Leptin/Adiponectin levels between controls (0.6±0.4) and cases (0.1±0.6), paired t-test, p=0.000.

The clinical questionnaires scores (CARS and ASQ) had no correlation with the leptin and adiponectin levels. Leptin levels for patients presented a positive correlation with weight (r=0.304, p=0.05), FM (r=0.390, p=0.02) and L/A ratio (r=0.368, p=0.019). The control group showed a positive correlation between leptin levels versus LDL-chol (r= 0.379, p= 0.016) and total-chol (r=-0.388, p= 0.013). Some weak to moderate correlations were found when testing correlations between leptin and adiponectin with clinical scores, anthropometric data and lipid profile (Table 3). Leptin levels presented positive correlations with weight and FM and were negatively correlated with HDL-chol in patients. In turn, adiponectin levels were negatively correlated with WC and AC in controls. L/A ratio correlated positively with weight and FM in ASD patients and with total-chol and LDL-chol in controls. Other anthropometric and lipid profile data, as well as the clinical scores (CARS and ASQ), had no correlation with the leptin and adiponectin levels and the L/A ratio. In addition, there was no correlation between leptin and adiponectin in both groups (r=-0.258, p=0.108 and r=-0.211, p=0.170 for controls and cases, respectively).

**Table 3.** Leptin and adiponectin correlations with anthropometrics (*weight, height, BMI, WC* and *AC*), body composition (*FM* and *FFM*), lipids (*HDL-chol, LDL-chol* and *total-chol*) and clinical (*CARS* and *ASQ*) variables ^†^

|  | Controls |  |  | ASD |  |  |
| --- | --- | --- | --- | --- | --- | --- |
|  | Leptin | Adiponectin | L/A | Leptin | Adiponectin | L/A |
| **Weight (kg)** | r= 0.083<br>p= 0.609 | r= -0.291<br>p= 0.069 | r= 0.168<br>p= 0.301 | r= 0.304<br>p= 0.057* | r= 0.025<br>p= 0.879 | r= 0.323<br>p= 0.042* |
| **Height (cm)** | r= -0.054<br>p= 0.739 | r= -0.201<br>p= 0.213 | r= -0.003<br>p= 0.987 | r= 0.269<br>p= 0.940 | r= 0.070<br>p= 0.669 | r= 0.243<br>p= 0.130 |
| **BMI (kg/m<sup>2</sup>)** | r= 0.094<br>p= 0.564 | r= -0.271<br>p= 0.091 | r= 0.175<br>p= 0.279 | r= 0.037<br>p= 0.822 | r= 0.138<br>p= 0.369 | r= -0.006<br>p= 0.970 |
| **WC (cm)** | r= 0.048<br>p= 0.767 | r= -0.361<br>p= 0.022* | r= 0.131<br>p= 0.419 | r= 0.289<br>p= 0.079 | r= -0.038<br>p= 0.822 | r= 0.279<br>p= 0.090 |
| **AC (cm)** | r= 0.136<br>p= 0.415 | r= -0.316<br>p= 0.050* | r= 0.226<br>p= 0.173 | r= -0.227<br>p= 0.159 | r= 0.227<br>p= 0.082 | r= -0.239<br>p= 0.138 |
| **FM (kg)** | r= 0.012<br>p= 0.942 | r= -0.258<br>p= 0.107 | r= 0.206<br>p= 0.203 | r= 0.390<br>p= 0.013* | r= -0.024<br>p= 0.883 | r= 0.368<br>p= 0.019* |
| **FFM (kg)** | r= 0.120<br>p= 0.462 | r= -0.299<br>p= 0.061 | r= 0.206<br>p= 0.203 | r= 0.239<br>p= 0.137 | r= 0.063<br>p= 0.701 | r= 0.224<br>p= 0.164 |
| **HDL-chol (mg/dL)** | r= -0.103<br>p= 0.526 | r= 0.184<br>p= 0.255 | r= -0.164<br>p= 0.312 | r= -0.464<br>p= 0.039* | r= -0.051<br>p= 0.753 | r= -0.087<br>p= 0.594 |
| **LDL-chol (mg/dL)** | r= 0.379<br>p= 0.016* | r= -0.015<br>p= 0.479 | r= 0.376<br>p= 0.075 | r= 0.251<br>p= 0.118 | r= -0.058<br>p= 0.721 | r= 0.251<br>p= 0.119 |
| **Total cholesterol (mg/dL)** | r= 0.388<br>p= 0.013* | r= 0.028<br>p= 0.960 | r= 0.405<br>p= 0.009* | r= 0.238<br>p= 0.140 | r= -0.083<br>p= 0.611 | r= 0.236<br>p= 0.142 |
| **CARS** | - | - | - | r= -0.258<br>p= 0.202 | r= 0.112<br>p= 0.588 | r= -0.220<br>p= 0.119 |
| **ASQ** | - | - | - | r= 0.012<br>p= 0.952 | r= -0.193<br>p= 0.326 | r= 0.034<br>p= 0.864 |

**AC:** Arm circumference; **ASQ:** Autism screening questionnaire; **BMI:** Body mass index; **CARS:** Childhood autism rating scale; **FFM:** Fat free mass; **FM:** Fat mass; **HDL:** High density lipoprotein cholesterol; **LDL:** Low density lipoprotein *cholesterol*; **WC:** Waist circumference.
<sup>†</sup>Pearson Correlation, \*Significant p-values

## 4. Discussion

Leptin and adiponectin have effects on the regulation of appetite and body composition. However, evidence of the relationship between these hormones and body composition in children is still limited (Dalskov et al., 2015) even in typically developing children. Even though investigations of their role in children with ASD are incipient, the nutritional aspects and eating difficulties that these patients may present are increasingly highlighted, often leading to inadequate nutritional statuses such as overweight and obesity (Castro et al., 2017; Gulati & Dubey, 2015). The main findings of our study report to a higher level of leptin and no changes in adiponectin levels in ASD children in comparison with controls, correlation of leptin with weight, FM and HDL-chol in ASD children, as well as higher FM and lower FFM in comparison with controls. In addition, L/A ratio was higher and correlated with weight and FM in patients. In the literature, adiponectin is negatively correlated with insulin resistance and obesity’ parameters like BMI (Maeda et al., 2002; Spranger et al., 2003; Weiss et al., 2004), body weight as well as with lean body mass and the lipid indicators LDL-chol and triglycerides (Lubkowska et al., 2015). In ASD patients, significant lower levels of adiponectin were observed when compared to healthy controls, suggesting a role of adiponectin in this pathology (Fujita-Shimizu et al., 2010). The same findings were reported in Rett Syndrome patients (Blardi et al., 2009). In contrast, in line with the present study, adiponectin levels did not differ between controls and young ASD patients (Blardi et al., 2010), even with a 1 year follow-up. Another paper found no differences between groups for adiponectin, but found a negative correlation with the severity of symptoms assessed by the Social Responsiveness Scale (Rodrigues et al., 2014). It is noteworthy that the analyzes of this study were performed controlling the effect of the BMI. Negative correlations were also found between adiponectin and impairment in social interaction in Fujita-Shimitzu et al. (2010) (Fujita-Shimizu et al., 2010), which, identified lower levels of adiponectin in ASD patients. None of these correlations was found in the present study, but the negative correlation of adiponectin with WC in controls could indicate that abdominal fat deposition is related to lower adiponectin levels. Furthermore, the negative correlation of adiponectin and estimated muscular protein (trough AC) in controls of our study could be compared with the data from Dalskow et al (2015) (Dalskov et al., 2015), which describes an inverse association between adiponectin and an FFM-index.

High serum leptin levels are widely discussed as a biomarker of adiposity in youth and adults (Ambroszkiewicz et al., 2017; Hamnvik et al., 2011; Willers et al., 2015).It has long been suggested that not only the consequences but also the etiology of obesity occurs through hyperleptinemia and leptin resistance (Zhang & Scarpace, 2006). Besides the present study, other authors have also reported higher leptin levels in ASD patients when compared to healthy individuals (Ashwood et al., 2008; Blardi et al., 2010; Rodrigues et al., 2014). Higher leptin levels were also reported for Rett Syndrome patients (Blardi et al., 2009). In our study, despite no different BMI means between patients and controls, the substantial higher body fat and the correlation with leptin levels in patients corroborate previous data of body composition of ASD children and adolescents found by our research group (Castro et al., 2017), and also data regarding the association between both in typically developing children and adolescents (Bundy, 2011; Li et al., 2016). Blardi et al. (2010)(Blardi et al., 2010), however, found no association between BMI and leptin levels, neither in ASD cases nor in controls, when analyzing this parameter, hypothesizing that since there were no obese in their sample, leptin might cooperate in clinical manifestations other than weight balance. In a specific overweight/obese heart failure patients study, leptin levels were not associated with obesity, however they were increased as the severity of obesity was greater (Motie et al., 2014). One must consider that BMI not necessarily reflects adiposity, emphasizing the importance of assessments of body composition. A recent meta-analysis, which analyzed the diagnostic performance of BMI to identify obesity as defined by body adiposity in children and adolescents, shows that BMI has high specificity but low sensitivity to detect the excess of adiposity, and fails to identify over a quarter of children with excess body fat percentage (Javed et al., 2015). In this sense, an important and broad longitudinal study that sought to evaluate the role of leptin in 8-11 years typically developing children found that baseline leptin was positively associated with an FM-index, supporting our cross-sectional data for ASD children, that leptin is produced in proportion to FM (Dalskov et al., 2015). However, our results regarding the absence of correlations between FFM and leptin in both groups were also reported in healthy children (Dalskov et al., 2015). The last cited study also points to a role of leptin as a negative predictor of subsequent gain in FFM, at least in girls; the authors consider that this finding, associated with the inverse relation with FM, may reflect preserved leptin sensitivity in their predominantly normal weight sample (Dalskov et al., 2015). The literature has a limited number of reports on body composition for ASD population (Castro et al., 2017) and, to our knowledge, none seeking the association of adipokines with body composition parameters, which limits the possibility of comparison with our results and points to important gaps to be explored in this issue.

Another element to be discussed is the higher L/A ratio in ASD patients compared to controls in this study. Whereas leptin is recognized as a proinflammatory cytokine, adiponectin downregulates the expression and release of many proinflammatory immune mediators (López-Jaramillo et al., 2014). Considering that adiponectin was negatively correlated with body weight (Lubkowska et al., 2015) and leptin was associated with FM (Dalskov et al., 2015), one can suppose that our data regarding the positive correlations of L/A ratio with weight and FM in patients could be related much more to higher leptin levels than to the L/A ratio, since adiponectin levels were similar between groups. In addition, there was no inverse correlation between leptin and adiponectin in our study.

The lipid profile and a possible relation with leptin levels was also investigated in our study. Jois et al. (2015) (Jois et al., 2015) report that in pre-pubertal children, leptin was a predictive variable for HDL-chol in males, and was related to insulin and lipid profile-namely HDL-chol, apoliprotein-A1 (apo-A1) and triglycerides - especially when leptin values are high (Jois et al., 2015). In the referred study, children in the highest terciles of leptin concentration had significantly lower values of HDL-chol and apo-A1and significantly higher triglyceride values than children in lower terciles (Jois et al., 2015). In another study, non-obese children in the highest quartile of L/A ratio demonstrated the lowest HDL when compared to lower ratio quartiles, whereas adiponectin levels were positively associated with HDL (Stakos et al., 2014). In the Fragile X Syndrome (FXS), a genetic disorder linked to ASD, lower levels of triglycerides, HDL, LDL, and total-chol were also described for patients when compared to controls (Çaku et al., 2017). In our sample, though there was no difference between groups in the lipid profile and despite the higher level of leptin in the patients, there was an inverse correlation between this and HDL-chol concentrations in the ASD group, whereas the L/A did not reach statistical significance. A reasonable explanation could be derived from the previously mentioned study findings (Jois et al., 2015), which reflect the relationship between leptin and lipid concentrations for healthy 6-8 years children with higher levels of leptin. Since these authors did not found a linear association between leptin and a worse lipid profile, they suggest a leptin resistant specific link between leptin and adverse lipid profile may occur (Jois et al., 2015).

Despite the positive correlation detected between leptin (and also the L/A ratio) versus total and LDL-chol in the control group, these findings did not appear to represent a distinct worse lipid profile between the groups, as can be seen in Table 3.

To avoid bias, we chose to include only participants without chronic use of medication in this study, since habitual drug use in patients with ASD may influence parameters related to the metabolism (Goel, Hong, Findling, & Ji, 2018; Sukasem et al., 2018).

Summarizing, this cross-sectional controlled study points to higher levels of leptin and no changes of adiponectin levels in ASD children in comparison with typically developing children. The novel finding of the positive correlation of leptin and FM in these patients supports its role as a marker of adiposity in ASD children, which is reiterated by the higher L/A ratio and its correlation with FM in patients. Inverse correlation of leptin with HDL-chol could be related to higher adiposity only in patients. These findings, despite contributing with important data, must be thoroughly explored and corroborated by additional studies. Besides, the complex regulation of body composition during childhood makes data related to hormonal aspects remarkably scarce in this specific population. Furthermore, additional longitudinal studies that are underway in our research group may increase the knowledge about the role of this adipokine in the body composition and lipid profile of children with ASD.

## Acknowledges

The authors would like to thank all the families who agreed to participate in the study the staff of Unit of Molecular and Protein Analysis (Experimental Research Center), Hospital de Clínicas de Porto Alegre, Porto Alegre, Brazil. The study was funded by Fundo de Incentivo à Pesquisa e Eventos-Hospital de Clínicas de Porto Alegre (FIPE-HCPA) (Grant Number: 16-0464). Public Brazilian agencies were neither involved in the study design and protocol, collection, analysis, and interpretation of data, in the writing of the report, nor in the decision to submit the paper for publication. KC is supported by Coordenação de Aperfeiçoamento de Pessoal de Nível Superior (CAPES). LSF was supported by Fundação de Amparo à Pesquisa do Estado do Rio Grande do Sul (FAPERGS).

Disclosure of interest, the authors report no conflict of interest.

